# FlashPCA2: principal component analysis of biobank-scale genotype datasets

**DOI:** 10.1101/094714

**Authors:** Gad Abraham, Yixuan Qiu, Michael Inouye

## Abstract

**Motivation:** Principal component analysis (PCA) is a crucial step in quality control of genomic data and a common approach for understanding population genetic structure. With the advent of large genotyping studies involving hundreds of thousands of individuals, standard approaches are no longer computationally feasible. We present FlashPCA2, a tool that can perform PCA on 1 million individuals faster than competing approaches, while requiring substantially less memory.

**Availability:** https://github.com/gabraham/ashpca

## 1 Introduction

Principal component analysis (PCA) of genotypes is an established approach for detecting and adjusting for population stratification and technical artefcats in genome-wide association studies (GWAS) and similar genomic analyses (Patterson *et al*., 2006; Novembre and Stephens, 2008; Price *et al*., 2010; Galinsky *et al*., 2016). The widely-used smartpca (EIGENSOFT) implementation has proven useful but it relies on two computationally-expensive steps: (i) computing the genetic relatedness matrix (GRM) 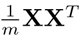 (**X** is the *n* × *m* matrix of standardised genotypes for *n* individuals and *m* single nucleotide polymorphisms, SNPs), and (ii) eigen-decomposition of the GRM. While this approach is effective for relatively small datasets (up to several thousand individuals), it becomes infeasible both in terms of memory requirements and computation time for larger datasets (𝒪(*mn*^2^) and 𝒪(*n*^3^), respectively).

Since most genomic analyses involving PCA only make use of the top 10–20 or so prinicpal components, alternative approaches that perform a partial decomposition have been proposed, including FlashPCA (Abraham and Inouye, 2014) and FastPCA (Galinsky *et al*., 2016). These tools have enabled analyses of far-larger datasets than would be practical otherwise. However, as demonstrated below, a downside of these algorithms is that they may not always converge rapidly to the solution or have substantial memory requirements. These shortcomings will be particularly challenging when analysing large datasets that are now becoming available, such as the UK Biobank (Sudlow *et al*., 2015) (*n* = 500,000 individuals) or the Precision Medicine Initiative (PMI) (Collins and Varmus, 2015), which intends to genotype 1 million individuals in the coming years.

Here we present FlashPCA2, which outperforms existing tools in terms of computation time on large datasets (*n* = 1,000,000 individuals and 100,000 SNPs), while utilising bounded memory and maintaining high accuracy for the top eigenvalues/eigenvectors. FlashPCA2 is implemented in C++ (based on the Eigen numerical library, http://eigen.tuxfamily.org), and relies on the Implicitly Restarted Arnoldi Method method as implemented in the C++ library Spectra (http://yixuan.cos.name/spectra).

## 2 Methods

Key to performance is the fact that the Arnoldi iterations only rely on vector-matrix multiplications with the genotype matrix **X**. FlashPCA2 employs a blockwise approach where a suitably-sized subset of the matrix is loaded into memory at one time. Other computational gains stem from precomputing the SNP-wise mean 2*p_j_* and standard deviation 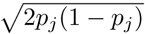, where *p_j_* is the minor allele frequency (MAF) for the *j*th SNP, only once and performing a lookup of these values in subsequent passes. In addition, in our experiments (below) the Arnoldi method exhibited better convergence than the algorithm of FlashPCA1, which can get stuck in local minima for before converging to the final estimates.

We used HAPGEN2 (Su *et al*., 2011), together with the 1000 Genomes 2009 CEU haplotypes (1000 Genomes Project Consortium, 2015) to simulate genotypes on chromosome 1 (600k SNPs) for up to 1,000,000 individuals. We used PLINK 1.9 (Chang *et al*., 2015) to thin the SNPs by linkage-disequilibrium down to 104,531 SNPs, making the SNPs approximately independent (Patterson *et al*., 2006). We first characterised the peformance of FlashPCA2 as a function of memory allocated to it (Supplementary Figure 1). Best performance was obtained when the data could be loaded into RAM fully, however, this strategy is not practical for large datasets, and we chose to allow a total of 2GiB RAM as a good compromise in terms of performance.

Next, we compared FlashPCA2 with FastPCA, FlashPCA1, and PLINK 1.9, examining wall run time, memory usage, and the error in the decomposition as a function of the number of individuals *n*, using *K* = 20 dimensions for FlashPCA and FastPCA. The error was defined as

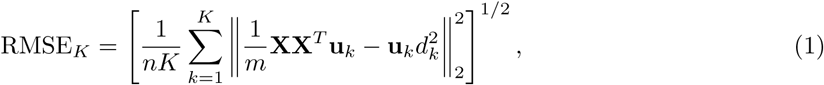

where ‖·‖_2_ is the Euclidean norm, **u**_*k*_ is the *k*th left singular vector of the GRM and *d_k_* is the *k*th singular value.

As Figure 1 shows, FlashPCA2 was the fastest, followed by FastPCA (2.6× slower), FlashPCA1 (8.9× slower), and finally PLINK (316× slower) (see Supplementary Figure 2 for run time as a function of the number of SNPs). Note that some FlashPCA1 and PLINK analyses could not be run due to the large memory requirements. FastPCA used up to 44GB RAM for the largest analysis (n=1,000,000), whereas FlashPCA2 used only 2GB RAM for all analyses. FlashPCA2 matched the accuracy of PLINK (full eigen-decomposition), followed closely by FlashPCA1, whereas FastPCA had RMSE 3–4 orders of magnitude higher (see Supplementary Figure 3 for results of up to 100,000 individuals).

**Figure 1:**
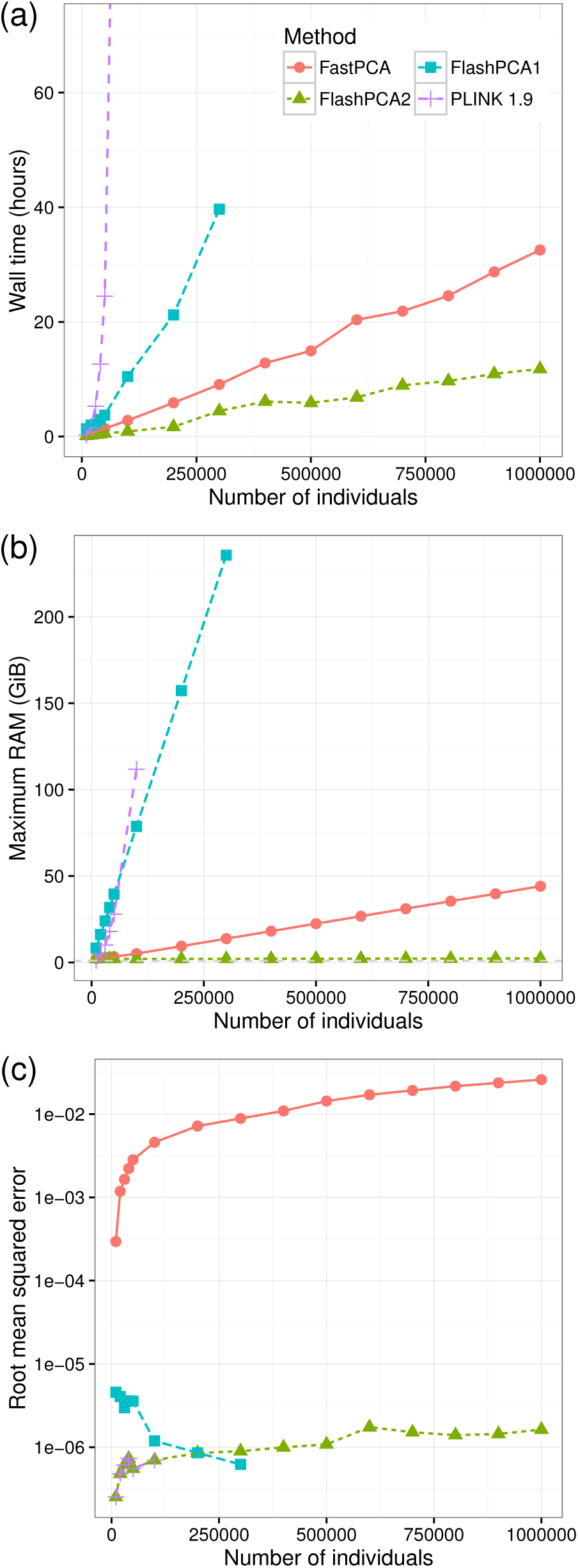
Performance on the HAPGEN2 chromosome 1 simulated datasets (104,531 SNPs), as a function of sample size. (a) Wall time (hours, average over 3 runs), truncated at 72h; (b) memory usage (GiB); (c) accuracy of the rank *K* = 20 decomposition (Equation 1).

## 3 Conclusion

FlashPCA2 enables scalable and accurate PCA of large genotype datasets, using small amounts of memory (2GiB for 1,000,000 individuals and 100,000 SNPs in <12 hours, single core), making it feasible to run such analyses on a standard personal computer, all within the **R** environment.

## Funding

GA was supported by a National Health and Medical Research Council Early Career Fellowship (NHMRC) no. 1090462. MI was supported by an NHMRC and Australian Heart Foundation Career Development Fellowship no. 1061435.

